# Plug-and-Play 3D localization microscopy

**DOI:** 10.64898/2026.07.29.741476

**Authors:** Ori Refael Cohen, Dafei Xiao, Reut Orange Kedem, Onit Alalouf, Prakash Joshi, Yuya Nakatani, Anna-Karin Gustavsson, Yoav Shechtman

## Abstract

Point-spread-function engineering by depth-encoding phase masks enables volumetric super-resolution imaging by 3D single-molecule localization microscopy (SMLM) but usually requires cumbersome relay optics. We demonstrate simple 3D SMLM implementation by phase mask insertion directly into the infinity space of a commercial microscope, along with appropriate computational compensation for field-dependence. The entire inserted component, including the phase mask, is 3D printing-based.

## Main

Single-molecule localization microscopy (SMLM)^1^ enables nanoscale imaging by precise determination of individual emitter positions^2–4^ and can be extended to three dimensions by encoding depth in the point spread function (PSF), i.e. PSF engineering ^5–7^. PSF engineering is implemented by shaping the wavefront on the microscope’s detection side, using spatial light modulators or diffractive optical elements (DOE). This typically necessitates significant modification to the optical setup by additional relay optics to access a plane conjugate to the back focal plane (BFP), leading to cumbersome alignment and a system-specific integration process^8,9^. Overall, these requirements limit the use of PSF engineering for 3D SMLM.

Here we introduce a plug-and-play (PnP) approach to PSF engineering in which a phase mask is inserted directly into the infinity space of a commercial inverted microscope, without any additional modification to the optical path. The PnP PSF-Engineering (PnP-PSFE) component consists of a wavefront-shaping optical element, i.e. a phase mask, mounted on a mechanical adapter compatible with an existing microscope slot (**Fig. 1**a). The entire component, including the Tetrapod phase mask demonstrated here^10^, is 3D-printing based^11^ fabricated as described in Supplementary Note 1. Unlike conventional implementations, the mask is not placed at a relayed BFP but directly in the microscope infinity space, between the objective and tube lens (**Fig. 1**b). Fluorescent bead stacks (Tetraspeck microspheres) confirm that the device generates depth-dependent Tetrapod-like PSFs over the field of view (**Fig. 1**c,d).

**Fig. 1:**
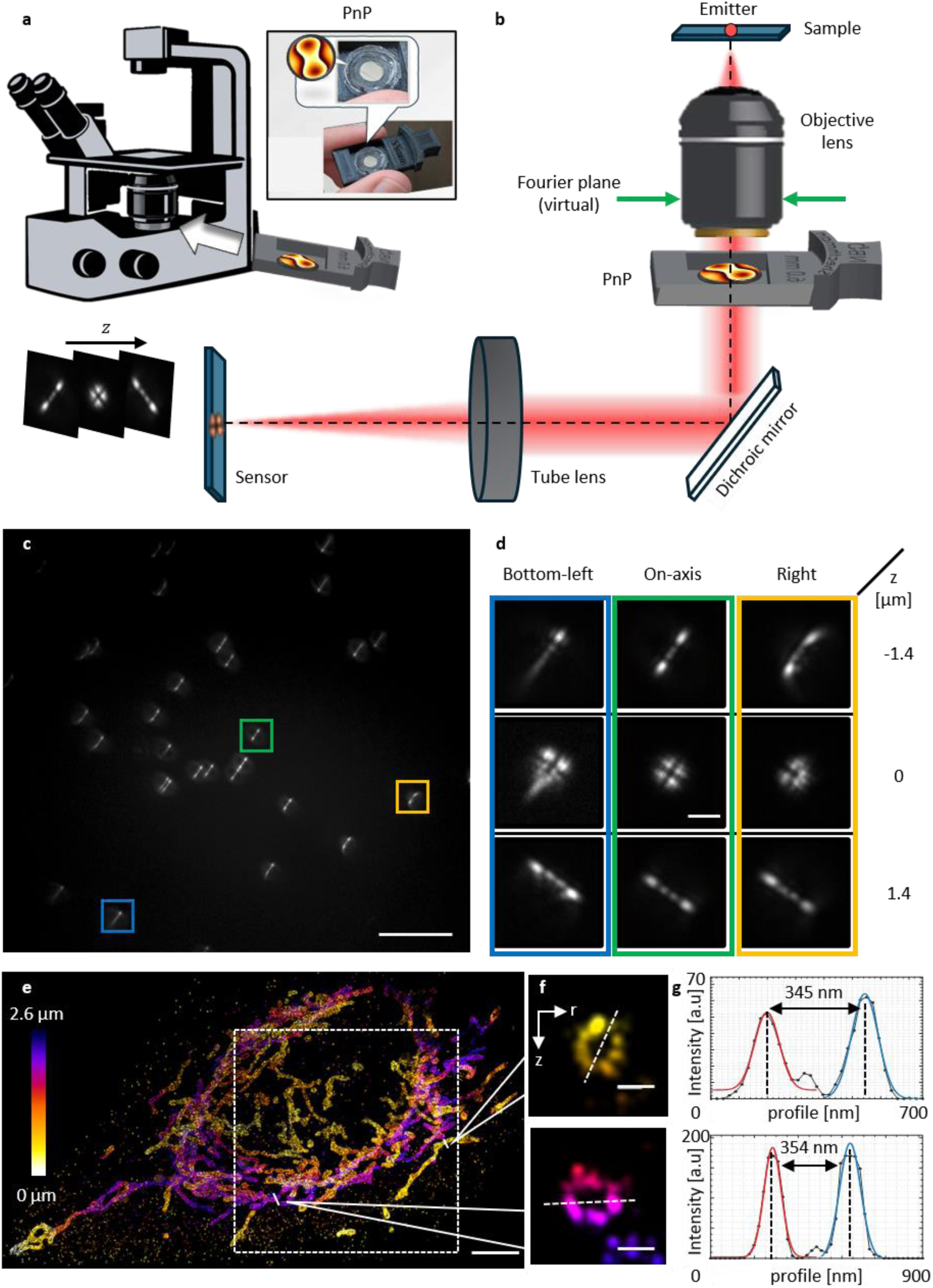
Plug-and-Play optical setup and near-axis 3D DNA-PAINT. **a**. PnP-PSFE module inserted into an inverted microscope. **b**. Detection path consists of a standard microscope imaging path, with the PnP component in the infinity space, modulating the phase to encode depth in the PSF shape. **c**. Fluorescent beads at ∼-1.4 μm defocus from focal plane, imaged with the PnP component inserted. A bead near the optical axis is marked in a green box, while beads at left-bottom side and right side relative to optical axis are marked in blue and orange, respectively. **d**. PSFs of the marked beads at −1.4, 0 and 1.4 μm defocus marked with corresponding box colors in **c**, shown in the top, middle and bottom rows respectively, exhibiting clear FOV-dependence. **e.** Near-axis 3D DNA-PAINT reconstruction of mitochondria, acquired with the PnP component and reconstructed using AutoDS3D. Color indicates axial position. Dashed box marks the ROI used for FRC analysis. **f.** (r)-(z) cross-sections along the solid lines in **e**. **g.** Profiles extracted along the dashed lines in **f** (in black), fitted with two Gaussians (left fit in red and right fit in blue); distances indicate fitted peak-to-peak separation of the hollow structure showing 345 nm (top) and 354 nm (bottom). Scale bars: 20 µm (**c**), 2 µm (**d**), 10 µm (**e**), and 0.25 µm (**f**).

Using the PnP component, we first demonstrated 3D DNA-PAINT imaging of mitochondria in a small region near the optical axis, where field-dependent PSF variation is minimal. The mask was inserted in a Nikon Eclipse Ti2 microscope DIC slot as in **Fig. 1**. Reconstruction using AutoDS3D^12^, a localization neural network, recovered 3D mitochondrial structures with 61 nm lateral resolution by Fourier ring correlation (FRC) analysis and enabled clear visualization of the hollow structure in r-z cross-sections (**Fig. 1**e,f).

While near the optical axis the measured PSF closely resembles the designed pattern, off-axis beads retain axial encoding but exhibit expected field-dependent distortions. This originates from the out-of BFP placement of the phase mask^13^, which leads to different regions of the mask being illuminated as a function of field-of-view (FOV) position (Fig. 2a). To characterize FOV dependence, which is necessary for downstream analysis of large-FOV imaging, we developed a numerical displaced-mask model, derived fully in Supplementary Note 2, in which the pupil field is propagated to the mask plane, modulated by the DOE and then mapped to the image plane. Notably, as part of the overall FOV-dependent PSF distortion, the displacement from BFP also produces apparent lateral displacement of the engineered PSF (Supplementary Note 3), which is modelled as well. In addition, signal-to-noise ratio (SNR) in the periphery of the FOV is reduced due to the aperture clipping and illumination losses, as discussed in Supplementary Note 4. A direct comparison with conventional Fourier-plane mask placement confirms that this field dependence is general to the displaced PnP configuration rather than to a specific phase mask design (Supplementary Note 5).

**Fig. 2:**
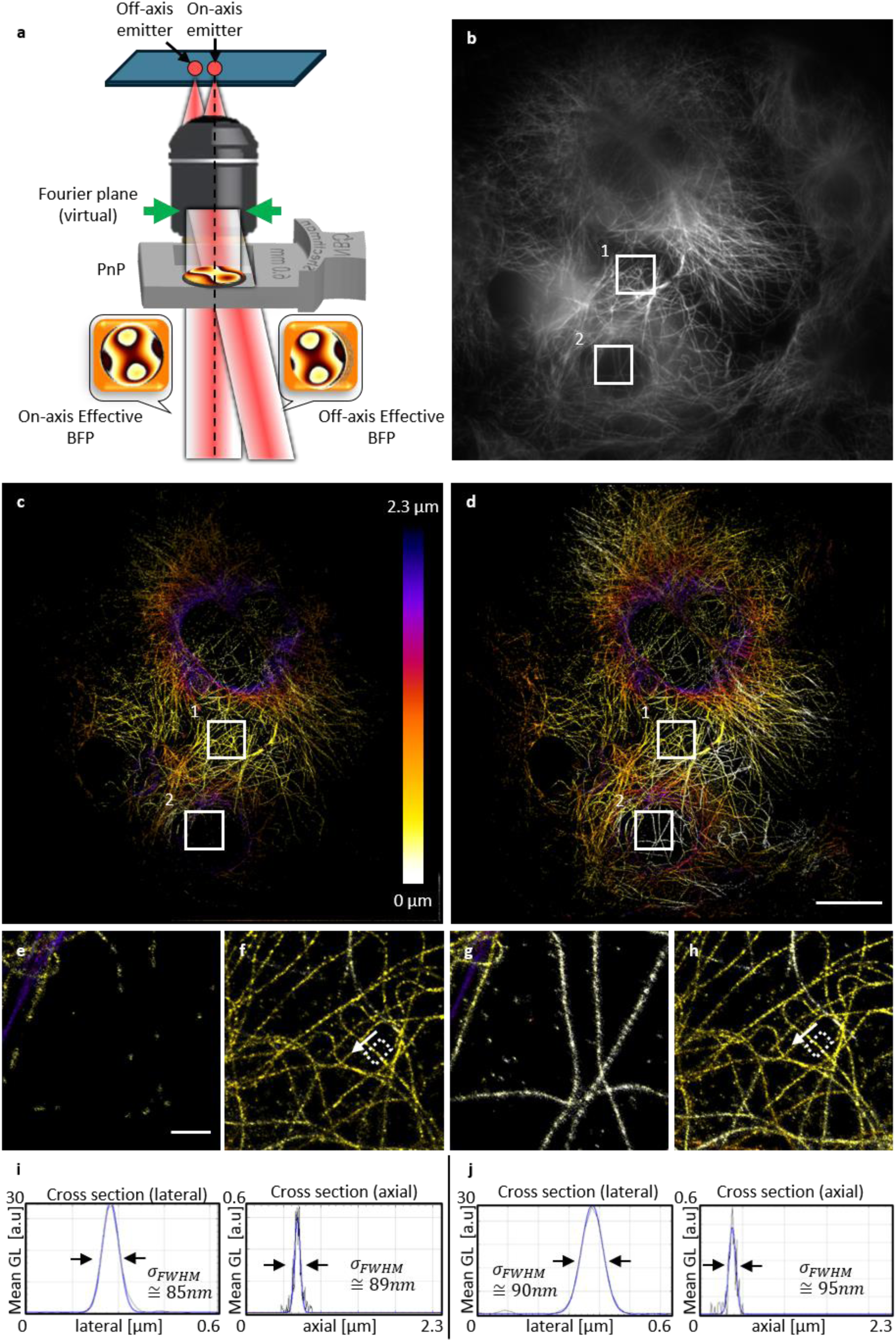
Large-FOV 3D dSTORM reconstruction of microtubules. **a**. Schematic illustration of the optical configuration: displacing the phase mask from the conjugate Fourier plane results in an apparent shift of the BFP for off-axis emitters, inducing spatially varying PSF distortions. **b.** Diffraction-limited widefield reference image of microtubules labelled with Alexa Fluor 647. **c**. Super-resolution dSTORM reconstruction processed with a standard shift-invariant localization algorithm (AutoDS3D), using an on-axis emitter for phase retrieval calibration; note the significant resolution degradation and structural blurring at the field periphery. **d**. Super-resolution reconstruction using the proposed field-dependent reconstruction, demonstrating restored structural integrity across the entire imaging area. **e-h.** 5× magnified views of the regions indicated by the solid white boxes (ROI 1 and ROI 2) in **b**-**d**, of the standard **e**,**f** and proposed **g**,**h** methods. **i**,**j.** Detailed comparison of a specific microtubule feature of the standard **i** and proposed **j** methods, indicated by the dashed white boxes in **f** and **h**, plotted as mean gray levels [a.u.] from ThunderSTORM-rendered images. Left plots: lateral cross-sections along the indicated arrows over a 100 nm slice (averaged perpendicular to the arrow direction). Right plots: ROI-averaged axial intensity profiles extracted from the dashed white boxes. For each 10 nm axial slice, the mean gray value within the box was calculated and plotted as a function of axial position. Scale bars: 20 μm (**b**–**d**) and 2 μm (**e**-**h**).

To improve downstream analysis of 3D emitter localization by deep-learning^7,12^, we incorporated a simple FOV-dependence calibration procedure consisting of measuring the z-stack of a few beads over the FOV, and fitting to our model. The complete workflow for calibration, synthetic training-data generation, network inference and localization are described in Supplementary Note 6. This allows us to generate training data that matches the measured PSF variation across the sensor, which is then used to train a FOV-dependent 3D localization neural net, based on Xiao et al.^14^. The displaced-mask phase-retrieval procedure is described in Supplementary Note 7.

We next tested the PnP component in large-FOV 3D dSTORM imaging of Alexa Fluor 647-labeled microtubules, where the mask was inserted similarly to the Mitochondria experiment. Blinking data were reconstructed using two analysis strategies, for quantifying the contribution of our FOV-dependent PSF model. A standard AutoDS3D^14^ workflow, with PSF calibrated using an on-axis bead and assuming a shift-invariant PSF, performed well near the optical axis but produced substantial blurring and loss of microtubule definition toward the field periphery (**Fig. 2**c). In contrast, the field-dependent reconstruction recovered continuous microtubule structures across the imaging area (**Fig. 2**d).

As expected, the total number of detected localizations was significantly higher when using the FOV-dependent model, compared to using AutoDS3D, with the same intensity threshold, namely, 4.9 million vs. 2.6 million localizations, respectively. This is most apparent in the periphery of the FOV, i.e. region of interest (ROI) 2 in **Fig. 2**c,d,e,g. Near the optical axis, in ROI 1 in Fig. 2b-d, where SNR is highest and the on-axis calibration best represents the PSF, the number of detected localizations was roughly similar, with slight advantage for AutoDS3D, with 69,500 for field-dependent reconstruction and 80,000 for AutoDS3D. FRC analysis on ROI 1 in Fig. 2 yielded lateral resolutions of 89 nm for both standard and field-dependent reconstruction. Cross-sectional analysis of isolated microtubules gave apparent widths of approximately 85 nm (AutoDS3D) and 90 nm (field-dependent reconstruction) laterally and 89 nm (AutoDS3D) and 95 nm (field-dependent reconstruction) axially (**Fig. 2** i,j). The dependence of axial profile width and localization density on the detection threshold is analyzed in Supplementary Note 8.

In summary, we achieve 3D SMLM by inserting a 3D-printing based PnP PSF engineering component into an existing commercial inverted microscope. Together with our previous compact spectral-encoding implementation^15^ and additional adapter geometries (Supplementary Note 9), these results support a practical strategy for PnP optical encoding. An Olympus-compatible adapter and an objective-mounted configuration are shown in Supplementary Note 9. Accounting for FOV-dependent PSF aberrations was done by developing a PSF model suitable for this configuration and training a FOV-aware 3D reconstruction net. Future work should consider larger mask areas and displacement-aware design of the phase mask pattern to prevent SNR loss in the periphery of the FOV. Overall, PnP-PSFE makes 3D localization microscopy readily available to practically anyone with an inverted microscope, without requiring any optical expertise or physical system modification, beyond trivial adapter-insertion.

## Online Methods

### Plug-and-play phase-mask fabrication and mounting

Diffractive phase masks were fabricated using 3D printing combined with near-index-matched polymers^11^. Briefly, a phase-mask template was fabricated using a microscale 3D printer, replicated into a polymer layer, and combined with a second closely index-matched polymer layer. The final diffractive optical element was enclosed between flat glass substrates to provide optical-quality interfaces. The phase mask was mounted in a custom 3D-printed mechanical adapter designed for insertion into the DIC slot of a Nikon Eclipse Ti2 inverted microscope. Additional details on fabrication, mounting and alternative adapter geometries are provided in Supplementary Notes 1 and 9.

### Microscope setup

Mitochondria and single-bead calibration experiments were performed on a Nikon Eclipse Ti2 inverted microscope equipped with a 100×/1.49 NA oil-immersion objective (CFI SR HP Apo TIRF 100x, NA 1.49 OIL, WD 0.12 mm, FOV 22 mm)^16^. The microscope was equipped with 561 nm laser excitation (1 W, 2RU-VFL, MPB Communications), and fluorescence emission was separated from excitation light using a multiband dichroic mirror (Chroma, ZT405/488/561/640rpcv2-UF3_91032) and filtered using a second dichroic mirror (Chroma, T590lpxr), a notch filter (Chroma, ZET561NF), and a bandpass filter (Chroma, ET605/70). The microscope image plane was relayed to the camera using a 1:1 two-lens relay composed of two 80 mm focal length lenses. Images were acquired on a Prime 95B sCMOS camera (Teledyne Vision Solutions) with 11 µm physical pixel pitch, corresponding to an effective camera pixel size of 109 nm in the sample plane.

Microtubule dSTORM imaging and field-dependent bead calibration experiments were performed on a Nikon Eclipse Ti2 inverted microscope equipped with a 100×/1.45 NA HP Plan Apo oil-immersion objective. Excitation was provided by a 640 nm laser, and fluorescence emission was separated from excitation light using a multiband dichroic mirror (Chroma TRF 89902-NK). The microscope image plane was relayed to the camera using a 1:1 two-lens relay composed of two 200 mm focal length lenses. This relay also enabled the Fourier-plane comparison described in Supplementary Note 5. Images were acquired on a Prime 95B camera with 11 µm physical pixel pitch, corresponding to an effective camera pixel size of 110 nm in the sample plane.

For PnP measurements, the phase mask was inserted directly into the microscope infinity space through the DIC component slot.

### Optical configuration and bead calibration

Fluorescent bead stacks were acquired to characterize the depth-dependent PSF via phase retrieval. Bead z-stacks were acquired over axial range used for model calibration. For the mitochondria experiment, Tetraspeck microspheres (T7280) beads measurements were performed using 561 nm excitation.

For the microtubules experiment and field-dependent calibration, Tetraspeck microspheres (T7280) bead measurements were performed using 640 nm excitation. A few additional beads across the FOV were used, when one of the beads that exhibited a symmetrical PSF was defined as on-axis bead; Additional measurements comparing displaced BFP and Fourier-plane mask placement are shown in Supplementary Notes 3–5.

The excitation laser power and exposure time were adjusted to obtain high-SNR images across the FOV without saturation.

### Displaced-mask image-formation model and phase retrieval

The field-dependent PSF was modelled using a displaced-mask image-formation framework, developed in this work. In this model, the complex pupil field is propagated from the BFP to the mask plane using the angular spectrum method (ASM), multiplied by the mask transmittance, and propagated back to the BFP before final image-plane transformation via a fast Fourier transform (FFT). To enable efficient computation on a downsampled mask grid, the linear phase term associated with lateral emitter displacement was omitted from the pupil field and instead applied as an equivalent lateral shift of the propagated field at the mask plane. The model therefore accounts for the fact that off-axis emitters interact with laterally shifted regions of a finite mask, producing field-dependent PSF deformation and apparent lateral displacement. The full derivation is provided in Supplementary Note 2.

To calibrate the model, experimentally measured bead z-stacks were used for phase retrieval. One bead near the optical axis was used as a reference, while additional off-axis beads constrained the field-dependent behavior. The recovered phase mask and estimated mask displacement were then used as fixed inputs for synthetic training-data generation for a FOV-dependent version of AutoDS3D^12^. Details of the displaced- mask phase-retrieval procedure are provided in Supplementary Note 7.

### Mitochondria DNA-PAINT sample preparation and imaging

Human osteosarcoma cells (U-2 OS – HTB-96, ATCC) were cultured in cell culture media (Dulbecco’s modified Eagle’s medium (DMEM) with 10% fetal bovine serum (FBS) and 1 mM sodium pyruvate, all Gibco) at 37°C at 5% CO_2_ and 90% humidity (Thermo Scientific Heracell 150i CO2 Incubator, 51-032- 871, Fisher Scientific). The cells were seeded into 8-well chambered coverslips (Ibidi GmbH). After incubation in the chambers for at least 12 h, the cells were fixed with 4% paraformaldehyde (PFA, Electron Microscopy Sciences) in phosphate-buffered saline (PBS) for 10 min at 37°C and washed three times with PBS and quenched by incubation with 10 mM NH₄Cl (Sigma-Aldrich, 213330) in PBS for 10 min at room temperature (RT), followed by three washes with PBS. Cells were permeabilized with 0.1% saponin in PBS for 10 min at RT and washed three times with PBS. Samples were then blocked for 1 h at RT in blocking buffer consisting of PBS supplemented with 3% (w/v) bovine serum albumin (BSA, Sigma-Aldrich, A2058), 0.05 mg/mL sheared salmon sperm DNA (Invitrogen, 15632011), 0.02% Tween-20 (Promega, H5152), and 0.05% sodium azide (G-Biosciences, 786-299). DNA-PAINT docking strand-conjugated secondary nanobodies were prepared following the protocol described previously^17^. Rabbit anti-TOMM20 primary antibodies (Abcam, ab186735) were preincubated with the DNA-conjugated anti-rabbit secondary nanobodies for 1 h at RT in antibody incubation buffer containing 1% (w/v) BSA, 0.05 mg/mL sheared salmon sperm DNA, 0.02% Tween 20 and 0.05% sodium azide in PBS. The preformed antibody–nanobody complexes were subsequently diluted to the desired working concentration (1:200 for primary antibodies) in antibody incubation buffer and incubated with the samples for 1–2 h at RT (or overnight at 4°C). Following labeling, samples were washed three times with PBS (5 min each) prior to imaging.

For single-molecule DNA-PAINT imaging of TOMM20-labeled mitochondria, Cy3B-conjugated imager strands (Massive Photonics) were introduced at a final concentration of 1 nM in imaging buffer (Massive Photonics). Fluorescence was excited using a 561 nm laser with an intensity in the sample plane of ∼340 W/cm^2^. Image sequences were acquired with an exposure time of 100 ms per frame, and 70,000 frames were recorded for single-molecule localization and DNA-PAINT image reconstruction.

### Microtubules dSTORM sample preparation and imaging

22 × 22 mm, 170 µm cover glasses (Deckgläser, No.1.5H) were cleaned in an ultrasonic bath with 5% Contrad 70 (Decon) at 60°C for 30 min, then washed twice with double distilled water, incubated in ethanol absolute for 30 min, sterilized with filtered 70% ethanol for 30 min and dried in a biological cabinet.

COS7 cells at a concentration of 45,000 cells/ml in DMEM with 1 g/l D-glucose (Thermo Fisher, 11880028), supplemented with fetal bovine serum (Biological Industries, 04-007-1A), penicillin-streptomycin (Biological Industries, 03-031-1B) and glutamine (Sartorius, 03-020-1B), were grown for 24 h in a 6-well plate (Thermo Fisher, Nunclon Delta Surface) containing 6 ml of the cell suspension and the cleaned cover glasses, at 37°C, and 5% CO_2_. The cells were fixed with 4% PFA and 0.2% glutaraldehyde in PBS, pH 6.2, for 60 min, washed 5 times, and incubated in 0.3 M glycine/PBS solution for 10 min. The cover glasses were transferred into a clean 6-well plate and incubated in a blocking/permeabilization solution for 2 h (10% goat serum, 3% BSA, 2.2% glycine, and 0.1% Triton-X in PBS, filtered with 0.45 μm Millex PVDF filter unit). The cells were then immune-stained with 1:250 diluted anti-α tubulin-AF647 antibody (Abcam, ab190573) in the blocking/permeabilization buffer for 1.5 h, shaking, and washed 5 times with PBS.

For super-resolution imaging, a PDMS chamber (22 × 22 × 3 mm, with a 13 × 13 mm hole cut in the middle) was attached to the cover glass containing the fixed and stained COS7 cells to create a pool for the blinking buffer. Blinking buffer (50 mM cysteamine hydrochloride (Sigma, M6500), 20% sodium lactate solution (Sigma, L1375), and 3% OxyFluor (Sigma, SAE0059) in PBS, pH 8-8.5) was added and a cover glass was placed on top while ensuring minimal air bubbles.

dSTORM image sequences were acquired using 640 nm excitation at 82 mW measured at the sample plane, with an exposure time of 40 ms per frame, and 50,000 frames were recorded for each reconstruction. During acquisition, weak 405-nm activation was applied intermittently as needed, up to 1 mW at the sample plane, to maintain a suitable density of blinking emitters.

### Network training and localization

Synthetic field-dependent training images were generated using the calibrated field-dependent forward model. Training for both AutoDS3D and the field-dependent reconstruction was performed on 150×150 pixel image patches. For the field-dependent reconstruction, the global field position of each pixel in each patch was preserved so that the simulated PSF matched the global pixel coordinate on the camera. AutoDS3D was trained on 10,000 patches, whereas the field-dependent reconstruction was trained on 3,000 full frames, with each frame divided into 64 patches. The network output was a three-dimensional volume with an effective camera pixel size of 110 nm, an axial voxel size of 28 nm, an axial range of 2.3 µm and frame full field of view was 132×132 μm. Experimental frames were processed by the trained networks and converted to emitter coordinates by thresholding the network output at 30, followed by three-dimensional non-maximum suppression and sub-voxel localization. A density filter requiring at least 30 neighboring localizations within a radius of 250 nm was applied to both AutoDS3D and field-dependent localization lists. The entire workflow is described in Supplementary Note 6.

### Mitochondria visualization and resolution analysis

Mitochondria DNA-PAINT data shown in **Fig. 1**e,f were acquired with the PnP component inserted into the microscope DIC slot. The imaged region was selected close to the optical axis, where field-dependent PSF variation is small. Therefore, this dataset was reconstructed using the standard AutoDS3D workflow with a single near-axis PSF calibration. Localization lists were visualized using the ThunderSTORM plugin in ImageJ. The 3D reconstruction was rendered with axial color coding at 5× magnification, corresponding to a pixel size of 22 nm. For the lateral view in **Fig. 1**e, localizations were projected in groups using 100 nm axial bins and ThunderSTORM averaged shifted histogram rendering with the lateral shift parameter set to 3.

For the r-z cross-sections in left panel of **Fig. 1**f, rendered image stacks were generated using 22 nm axial slices and ThunderSTORM averaged-shifted-histogram rendering with lateral and axial shift parameters set to 3. Intensity profiles were extracted along the dashed lines in the r-z cross-sections. Each profile was fitted with two one-dimensional Gaussian functions, and the reported distance was calculated as the peak-to-peak separation between the fitted Gaussian centers. Lateral resolution was estimated by FRC from the x-y projection of the dashed ROI in **Fig. 1**e, using odd and even frames as statistically independent subsets and the 1/7 threshold criterion.

### Microtubule visualization and resolution analysis

Localization lists of the microtubules were visualized using the ThunderSTORM plugin in ImageJ. Full-FOV of reconstructed images were rendered without magnification, corresponding to a pixel size of 110 nm. Magnified regions of interest were rendered at 5× magnification, corresponding to a pixel size of 22 nm. For lateral views, localizations were grouped into 100 nm axial slices. For axial-profile analysis, rendered image stacks were generated using 10 nm axial slices, and axial profiles were extracted as the mean gray value within the indicated region of interest in each axial slice. ThunderSTORM lateral shift and axial shift parameters were used as part of averaged shifted histogram rendering and were kept identical between corresponding AutoDS3D and field-dependent reconstructions.

FRC was used to estimate lateral resolution in the selected region of interest indicated in the main text. Localizations were divided into two statistically independent subsets according to frame parity, with odd and even frames reconstructed separately into two super-resolution images. The two images were transformed into Fourier space, and the FRC curve was computed as the normalized correlation between corresponding Fourier rings. The reported resolution was taken as the inverse of the spatial frequency at which the curve crossed the 1/7 threshold. The effect of the detection threshold on localization density and apparent axial width is described in Supplementary Note 8.

### AI-assisted tools

Large language model tools, including ChatGPT, were used during manuscript preparation for language editing, structural suggestions and assistance with drafting or debugging analysis code. All text, code, analyses and conclusions were reviewed, edited and approved by the authors. ChatGPT was not used as an author and was not used to generate, alter or interpret experimental data or figures.

## Acknowledgements

We are grateful to Prof. Mike Heilemann and Alexandra Kaminer for their collaboration on method validation, Ofri Goldenberg for his effort on algorithm development and Amit Parizat for his help on holder design.

## Data availability

Representative raw datasets and design files are available on Zenodo. The mitochondria DNA-PAINT dataset, including 50 raw frames and bead calibration z-stack used for PSF calibration, is available at https://doi.org/10.5281/zenodo.21648990. The microtubule dSTORM dataset, including 50 raw frames, full FOV bead calibration z-stack and cropped stacks used for PSF calibration, is available at https://doi.org/10.5281/zenodo.21648914. STL design files for the Nikon holder, Olympus holder and Olympus lid are available at https://doi.org/10.5281/zenodo.21648812. Additional data is available from the corresponding author upon reasonable request.

## Code availability

Mitochondria DNA-PAINT data and the non-FOV-aware microtubule reconstruction were processed using AutoDS3D, which is publicly available at: https://github.com/alonsaguy/One-click-image-reconstruction-in-single-molecule-localization-microscopy-via-deep-learning. The code used for field-dependent simulation, training and reconstruction is available from the corresponding author upon request.

## Author contributions

O.R.C. fabricated the plug-and-play component, collected the dSTORM data, reconstructed the 3D datasets, developed the displaced-mask imaging model, implemented the FOV-dependent reconstruction pipeline, analyzed the data and wrote the manuscript. R.O.K. contributed to phase-mask fabrication and advised on experiments and FOV-dependent algorithm development. D.X. advised on AutoDS3D usage, experimental design, manuscript preparation and development of the FOV-dependent reconstruction algorithm. O.A. helped with dSTORM data preparation. J.P. and Y.N. contributed to DNA-PAINT sample preparation and data acquisition. A.-K.G. reviewed and edited the manuscript. Y.S. supervised the project, advised on experimental design, contributed to dSTORM data acquisition, FOV-dependent algorithm development, data analysis and manuscript writing. All authors reviewed, contributed to and approved the final manuscript.

## Competing interests

The authors declare no competing financial interest.

## Funding

Funding was received from the Israel Science Foundation (grant No. 1081/24) to Y.S. This work was supported by the National Institute of General Medical Sciences of the National Institutes of Health grant R35GM155365 and startup funds from the Cancer Prevention and Research Institute of Texas grant RR200025 to A.-K.G.

## Supporting information

### Supplementary Note 1 – Plug and play component fabrication

The masks in this work were fabricated using 3D printing combined with near-index matched polymers^1^. The axial dimension of the element was scaled by orders of magnitude relative to conventional fabrication approaches, such as photolithography, enabling fabrication with a commercial 3D printer while maintaining high optical quality. A phase-mask template was fabricated using a microscale 3D printer (Fabrica Giga 25vx high-resolution). A first polymer was cast onto the template, cured at 75 °C for several hours, and released to form a negative replica constituting the first layer. A second layer with a closely matched refractive index was then added. To provide optical-quality interfaces at both air-facing surfaces of the DOE, the structure was sandwiched between two commercial flat glass substrates. The final device therefore consisted of a flat phase element formed by two closely index-matched layers enclosed between glass substrates.

Building on our previous compact spectral-encoding implementation^2^, the phase mask was mounted and glued to a custom 3D-printed adapter designed for insertion into the dedicated slot of a Nikon Eclipse Ti2 inverted microscope. Since the adapter geometry can be easily redesigned, the method is readily transferable to other commercial microscope platforms. In this work, the DOE was placed in the slot intended for DIC components.

### Supplementary Note 2 – Physical model of a displaced phase mask

We modeled image formation as coherent propagation of the emitter’s complex optical field through three planes: the pupil plane (back focal plane), the phase-mask plane and the image plane.

In the scalar field theory, the field in the image plane *E_img_*(*x_img_*, *y_img_*|*x_o_*, *y_o_*, *z_o_*) due to a point source at position (*x_o_*, *y_o_*, *z_o_*) in the object plane is proportional to the Fourier transform of the field at the pupil plane (Fourier plane)^3^:

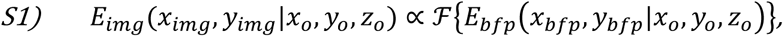

and the PSF, *I*, is proportional to the absolute value squared

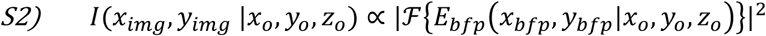

For an emitter at position (*x_o_*, *y_o_*, *z_o_*) in the object plane, the complex optical field at the back focal plane (BFP), *E*_0_*_bfp_*(*x_bfp_*, *y_bfp_*|*x_o_*, *y_o_*, *z_o_*), is determined by axial and lateral phase terms; the axial term depends on the emitter defocus, whereas the lateral term corresponds to the off-axis phase tilt of the wavefront. Therefore,

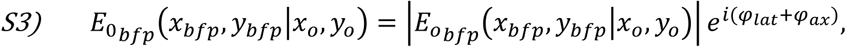

Whereas

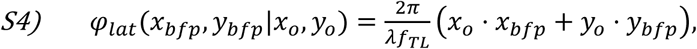

and the axial phase can be written as

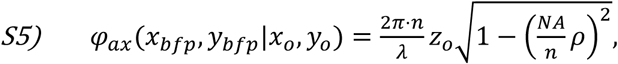

Whereas *n* is the refractive index of the immersion medium, *ρ* is the normalized pupil coordinate such that 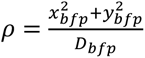 and *D_bfp_* is the pupil physical diameter of the objective lens^4^, 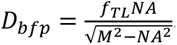

To model the effect of a phase mask located away from the nominal pupil plane, we propagate the complex field at the BFP in free space to the mask plane, using the angular spectrum method (ASM). Denoting the propagation distance by *d*, the field at the mask plane is

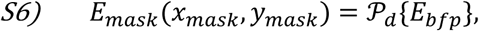

where P*_d_*{⋅} is the ASM propagation operator by distance *d* . The phase mask is then applied multiplicatively,

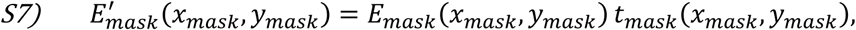

With 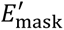 is the complex electric field at mask plane, after mask is applied, and

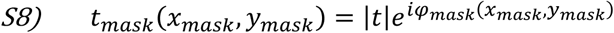

Is the mask complex transfer function, where in the case of DOE phase mask with Iris, |*t*| = 1 inside the radius and *φ_mask_*(*x_mask_*, *y_mask_*) is the phase modulation. The modulated field is propagated back to the pupil plane,

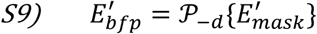

The image-plane field is given by the Fourier transform of the modulated pupil field,

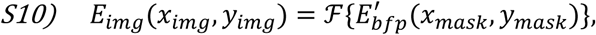

and the corresponding point spread function (PSF) is

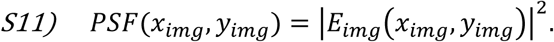

Thus, the phase mask modifies the pupil-plane wavefront, and after Fourier transformation this modulation is expressed as a reshaped PSF in the image plane.

From Eq. S6, and Eq. S9-S11, for an emitter at field position ***ρ*** and axial position *z*, the field-dependent PSF can be written as

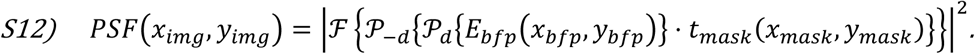

A key point in the off-axis case is that the lateral phase term is linear across the pupil. In Fourier optics, a linear phase corresponds to a lateral steering of the beam. Therefore, when the emitter is displaced from the optical axis by a distance (*x_o_*, *y_o_*), the field is no longer incident on the mask plane in the same manner as the on-axis field. Instead, the propagated wave is shifted laterally at the mask plane and effectively interacts with a different region of the phase mask. As a result, off-axis emitters do not experience a simple translated copy of the on-axis PSF; rather, they experience a position-dependent modulation determined by the local mask region sampled by the steered wavefront. This mechanism provides a natural physical description for field-dependent PSF variation.

After propagation, the simulated PSF is normalized to the photon count, mildly blurred to account for residual system blur, and cropped to the relevant camera region. The same physical forward model is used in phase retrieval, training-data generation, and localization, so that the experimentally calibrated mask and mask displacement are consistently incorporated throughout the workflow.

### Supplementary Note 3 – Apparent lateral displacement induced by off-pupil mask placement

Placing a finite phase mask away from a pupil-conjugate plane causes the mask to act not only as a phase element, but also as an aperture located at a non-pupil plane. For off-axis emitters, the beam footprint is laterally shifted at the mask plane and can therefore be asymmetrically modulated or partially clipped. As a result, the engineered PSF may appear laterally displaced relative to the actual emitter position.

To visualize this effect experimentally, we acquired images of the same fluorescent beads with and without the PnP phase mask. The unmasked bead image provides a reference for the true lateral bead position, whereas the masked z-stack shows the depth-encoded PSF produced by the displaced phase mask. As shown in **Fig. S3**a, the no-mask bead positions remain compact and focused, while the corresponding PnP PSFs are extended and field dependent. Magnified overlays of representative beads are shown in **Fig. S3**b. Near the optical axis, the engineered PSF remains approximately centered on the unmasked bead position throughout the axial scan. In contrast, off-axis beads show clear PSF deformation and apparent lateral displacement relative to the unmasked reference.

**Fig. S3:**
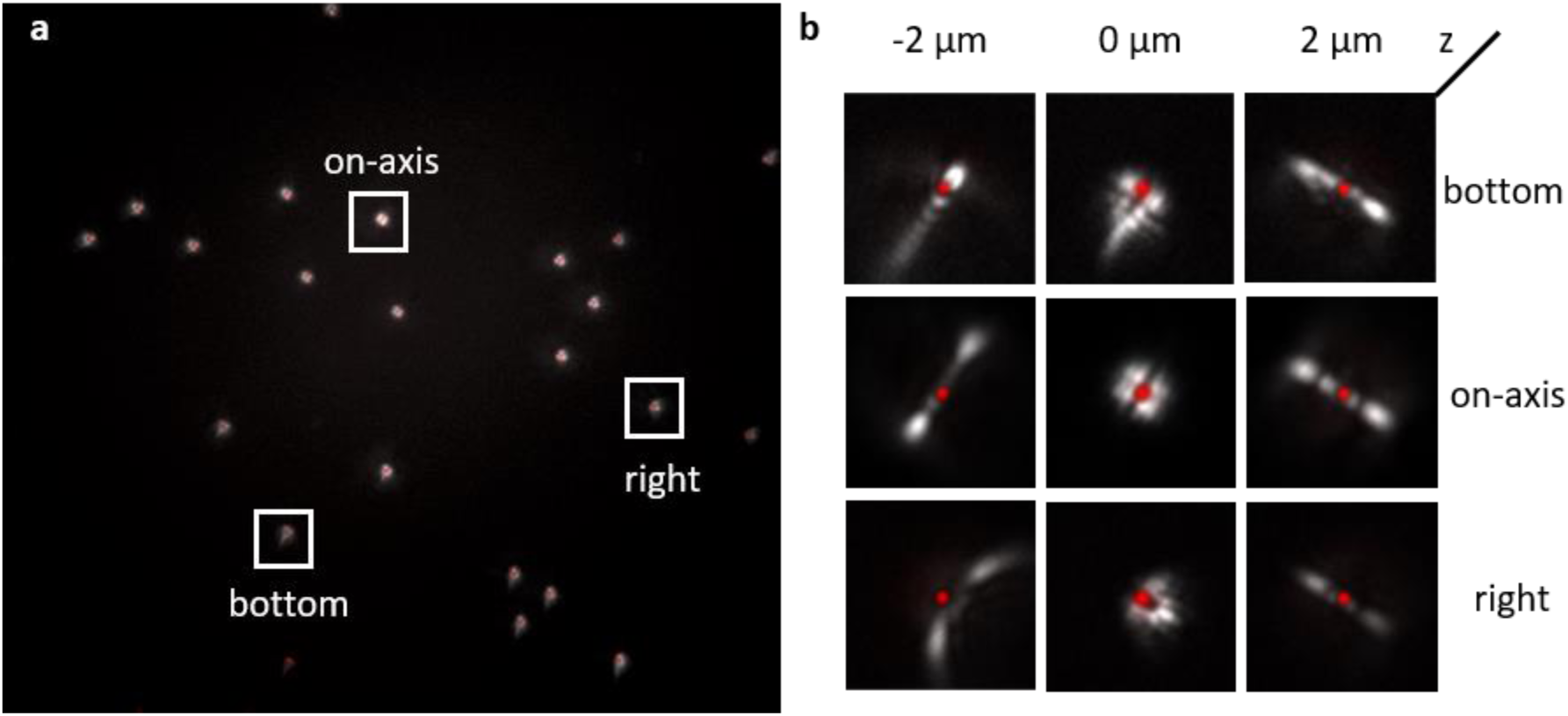
Field-dependent lateral displacement in the PnP configuration. Field-dependent lateral displacement in the PnP configuration. **a,** Full-field overlay of fluorescent beads imaged without the phase mask and with the PnP phase mask inserted. The red signal corresponds to the same beads imaged without the mask and provides a reference for their true lateral positions. The gray signal corresponds to the bead z-stack acquired with the PnP mask, showing the depth-encoded engineered PSFs. White boxes mark three representative beads located near the optical axis, at the bottom of the field of view and at the right side of the field of view. **b,** Magnified overlays of the boxed beads at selected axial positions. The top, middle and bottom rows correspond to the bottom, on-axis and right beads, respectively; the left, center and right columns correspond to different axial planes in the z-stack. Near the optical axis, the engineered PSF remains approximately centered on the red unmasked reference. Away from the optical axis, the PSF becomes deformed and laterally displaced relative to the true bead position, demonstrating the field-dependent lateral bias introduced by off-pupil mask placement. The unmasked images are all in the same focus plane and used as a reference point.

This apparent displacement can introduce lateral localization bias when the data are analyzed using a single shift-invariant PSF calibration. Because the engineered PSF shape also varies with defocus, the apparent displacement may change over the axial range, producing z-dependent lateral drift if field dependence is not considered. This observation further motivates field-dependent calibration and reconstruction, in which each emitter is modeled using the local PSF expected at its position in the field of view.

### Supplementary Note 4 – Signal-to-noise reduction induced by displaced mask placement

One limitation of the present plug-and-play configuration is reduced signal-to-noise (SNR) ratio compared with a conventional 4f implementation. Two effects contribute to this loss. First, fluorescence transmission through the DOE can be reduced by Fresnel reflection, scattering or absorption. Second, because the mask has a finite active aperture, displacement from the pupil plane causes off-axis emission to interact with a shifted region of the mask (see ray trace introduced in Fig. 2a in the main text). When the mask aperture is matched to the nominal back focal plane diameter, this can partially clip off-axis beams and produce vignetting-like signal loss.

Both effects can be mitigated in future implementations. Optical throughput can be improved by optimized fabrication and anti-reflection coatings. In addition, a larger active mask area can be designed to prevent clipping of off-axis beams across the intended field of view. These modifications should improve peripheral SNR while preserving the practical plug-and-play implementation.

### Supplementary Note 5 – Comparison with Fourier-plane mask placement

In conventional PSF engineering, a diffractive optical element is positioned at a plane conjugate to the objective pupil. This is commonly achieved using a 4f relay, in which two lenses separated by the sum of their focal lengths provide physical access to the Fourier plane. In this configuration, the phase mask modulates the pupil field in a field-independent manner, producing an approximately invariant engineered PSF over the measured field of view.

To compare this conventional geometry with the PnP configuration, we used a 1:1 detection relay and placed the phase mask either at the intermediate Fourier plane between the two 200 mm lenses or directly in the microscope DIC slot behind the objective. Full-field bead images acquired in the two configurations are shown in Fig. S4a. Magnified PSFs from representative field positions are shown in Fig. S4b, demonstrating that Fourier-plane mask placement produces an approximately field-invariant PSF, whereas direct PnP insertion produces the expected field-dependent deformation.

**Fig. S4:**
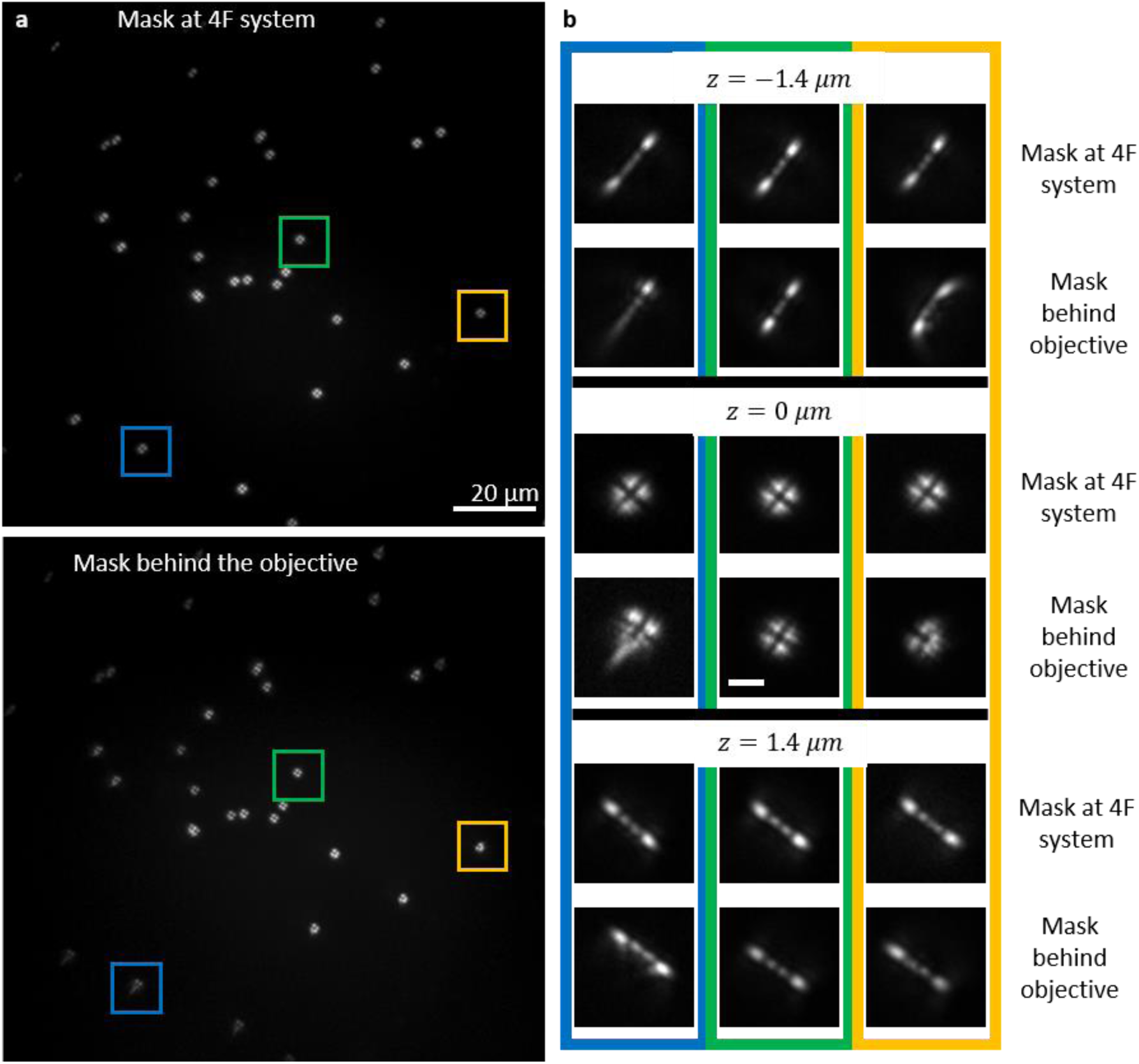
Comparison of PSF across the field of view for different mask placement configurations. Comparison between the Point Spread Functions (PSFs) when the phase mask (DOE) is placed in the conventional 4f-system Fourier plane (top full-field image) versus the proposed Plug-and-Play (PnP) configuration behind the objective (bottom full-field image). **a**, Large field-of-view (FOV) images showing multiple fluorescent beads, where specific regions are marked (colored boxes) corresponding to different field positions. **b**, Zoomed-in PSFs from these marked regions, displayed at selected axial positions (*z* = −1.4, 0 *and* 1.4 *μm*). As described in the supplementary text, positioning the mask at the true Fourier plane (4f system) results in a highly field-invariant PSF. In contrast, the off-plane PnP position (behind the objective) introduces field-dependent aberrations, leading to field-dependent distortions, particularly at the FOV periphery (compare left and right column zooms). Scale bars are 20 µm in the large images and 2 µm in the zoomed-in ROI.

### Supplementary Note 6 – Computational pipeline

The software pipeline was developed as a unified framework that bridges optical calibration, simulation-based training, and experimental inference. The process begins with optical calibration, where experimental bead z-stacks are analyzed via phase retrieval to characterize the system. This step estimates the effective phase mask and additional parameters, such as mask displacement and the nominal focal plane offset, enabling the forward model to accurately reproduce measured PSFs across the entire field of view.

Once calibrated, the forward model is employed to generate synthetic training data. By randomly sampling fluorophore positions throughout a 3D imaging volume and simulating emitter images with field-dependent PSFs, the system creates a realistic dataset that resembles experimental conditions. As demonstrated in **Fig. S5**, the simulation explicitly incorporates realistic noise statistics and background characteristics, ensuring that each generated frame paired with its 3D ground-truth coordinates trains the model for experimental data.

**Fig. S5:**
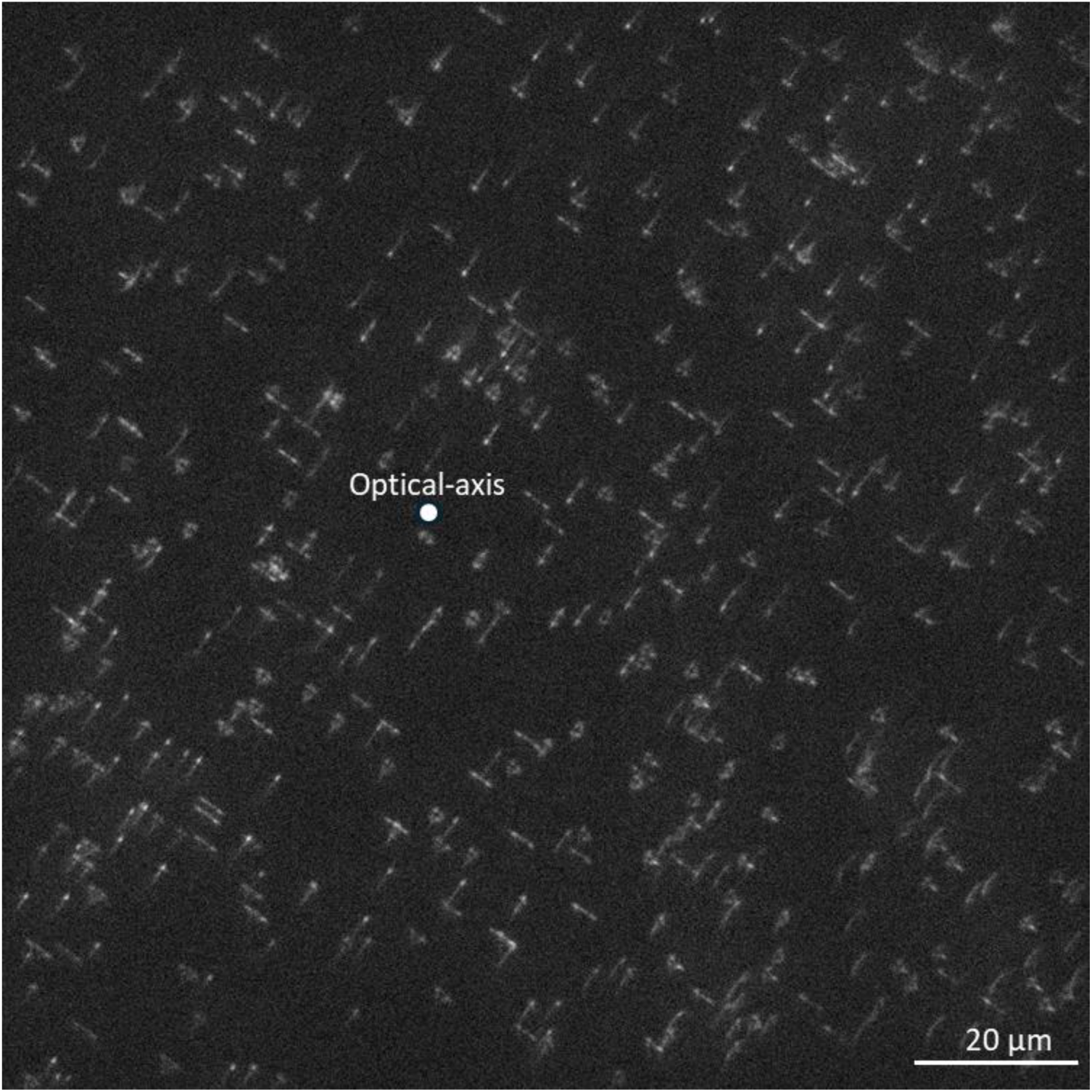
Synthetic training data generation. A representative simulated frame used for network training, showing a high density of emitters with spatially varying PSFs. The simulation incorporates field-dependent distortions and experimental noise levels to ensure high-fidelity localization during inference. The optical-axis point is similar to the experimental data, at pixel coordinates (477, 573) is marked by a white dot. Scale bar: 20 µm.

In the third stage, these simulated data were used to train a convolutional neural network to map a single input image into a three-dimensional emitter volume. The training procedure therefore relied entirely on model-based synthetic supervision, while explicitly incorporating the spatially varying PSF across the field of view.

In the final stage, experimental image sequences were processed by the trained network to produce a 3D output volume for each frame. These volumes were then converted into emitter coordinates by applying thresholding, non-maximum suppression, and local sub-voxel refinement. This approach is visualized in **Fig. S6**, which showcases the network’s ability to maintain localization consistency even in dense, low-SNR regions. For large fields of view, inference could be performed in a patch-wise manner while preserving each patch’s global field position, thereby maintaining consistency with the field-dependent training model.

Overall, this pipeline provided a unified framework in which the calibrated optical model informed both the generation of supervision for network training and the interpretation of experimental data during localization.

**Fig. S6:**
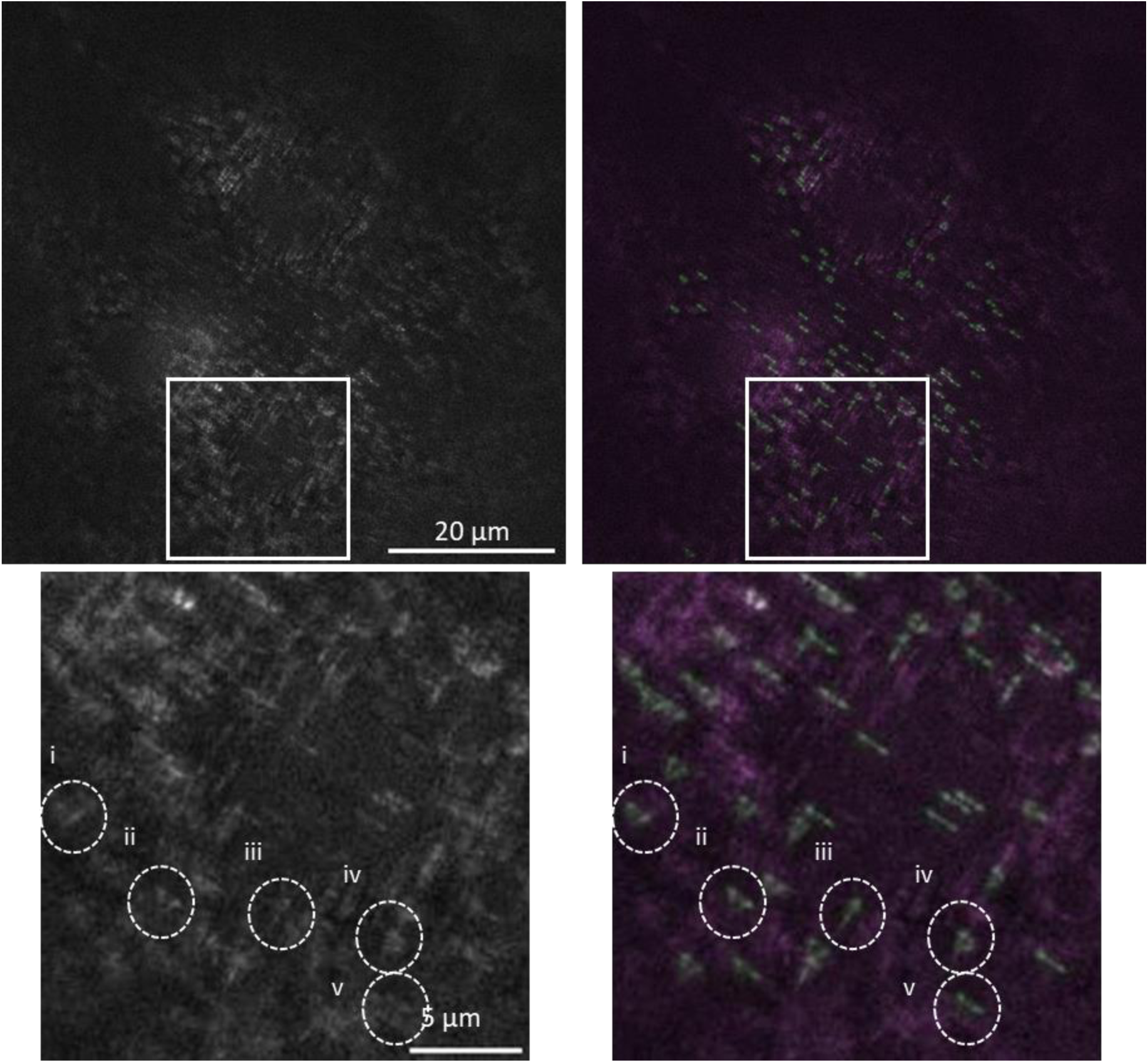
Experimental 3D localization considering FOV dependence. Demonstration of the computational pipeline’s performance on experimental data. Top row: Full FOV raw single frame acquisition in the left panel and the resulting 3D localizations (green), overlaid on the input frame in the right panel (purple) (right). Bottom row: Magnified views of the region indicated by the white box, illustrating the high-fidelity reconstruction of dense structures. The numbered circles (i–v) highlight specific emitters that were accurately localized despite significant local PSF distortions, PSF overlap and low SNR. These results illustrate that the calibrated forward model enables the neural network to handle field-dependent PSF variation in experimental data. Scale bars: 20 µm (top) and 5 µm (bottom).

### Supplementary Note 7 – Phase retrieval for displaced DOE

To calibrate the imaging model, we estimated the phase mask directly from experimentally measured bead z-stacks. The goal of this optimization step was to recover a forward model whose simulated PSFs match the measured PSFs, so that the same model could later be used for training-data generation and inference. Phase retrieval is a well-known approach in the field. However, it usually retrieves phase from a spatially invariant linear PSF generated from a bead. In our work, we implemented phase retrieval for a phase mask placed away from the pupil plane. The inputs are z-stacks from several beads across the FOV, and the outputs are the recovered phase mask and the estimated mask displacement from the pupil plane. Thus, by analyzing typically 3-8 beads, This allows the recovered model to reproduce the field-dependent PSFs generated by the displaced mask.

One bead near the optical axis was used as the reference bead, while additional off-axis beads constrained the field-dependent behavior of the model. For each bead, a z-stack was acquired over the relevant axial range. Background was first reduced and each slice was normalized to emphasize PSF shape rather than absolute intensity. The phase mask was then optimized by minimizing the discrepancy between measured and simulated PSFs produced by the forward model. Representative measured and simulated PSFs from beads at different field positions are shown in **Fig. S7**a, demonstrating that the calibrated model reproduces the main field-dependent PSF changes across the axial range.

The optimization was performed directly on the mask phase in the mask plane. At each iteration, the current phase mask was inserted into the imaging model, propagated to produce simulated PSFs for the measured bead positions, and compared to the corresponding experimental z-stack slices. The loss function was based on the agreement between simulated and measured PSFs, while allowing robustness to small residual alignment errors both laterally and axially. In particular, for off-axis beads, the comparison could include limited lateral alignment and small bead-specific axial offsets in order to reduce sensitivity to imperfect experimental bead centering and focus selection. This made the recovered mask less dependent on manual selection of the exact bead center or nominal focal slice.

Because off-axis emitters induce large pupil-plane phase slopes, direct optimization can become numerically unstable on a practical pupil-plane grid. Therefore, during phase retrieval we used the same method described above, in which the lateral phase was omitted and replaced by an equivalent lateral shift after propagation. This allowed stable simulation of field-dependent PSFs without increasing the pupil-plane sampling density. The corresponding effective pupil-plane phase experienced by emitters at different field positions is shown in **Fig. S7**b, illustrating how the displaced mask produces position-dependent modulation.

The phase mask was optimized jointly with the mask displacement parameter (d). After convergence, the recovered phase mask and mask displacement were used as fixed inputs for synthetic training-data generation. In this way, the simulated training PSFs inherited the experimentally calibrated optical behavior of the system, including the field dependence induced by the displaced mask.

During development of the modified phase-retrieval procedure, we found that the nominal-focal-plane (NFP) parameter strongly affected the agreement between simulated and measured PSFs in the displaced- mask configuration. While this parameter is typically assumed to be near NFP=0 for standard bead calibration, we found that setting NFP=-3 μm, yielded the best fit between our model and the experimental data. This adjustment accounts for the fact that a system with a mask placed behind the objective is not perfectly aligned. Specifically, it compensates for a small defocus from the nominal working distance, an offset that is generally negligible in traditional systems but becomes dominant when the mask is displaced from the pupil plane.

**Fig. S7:**
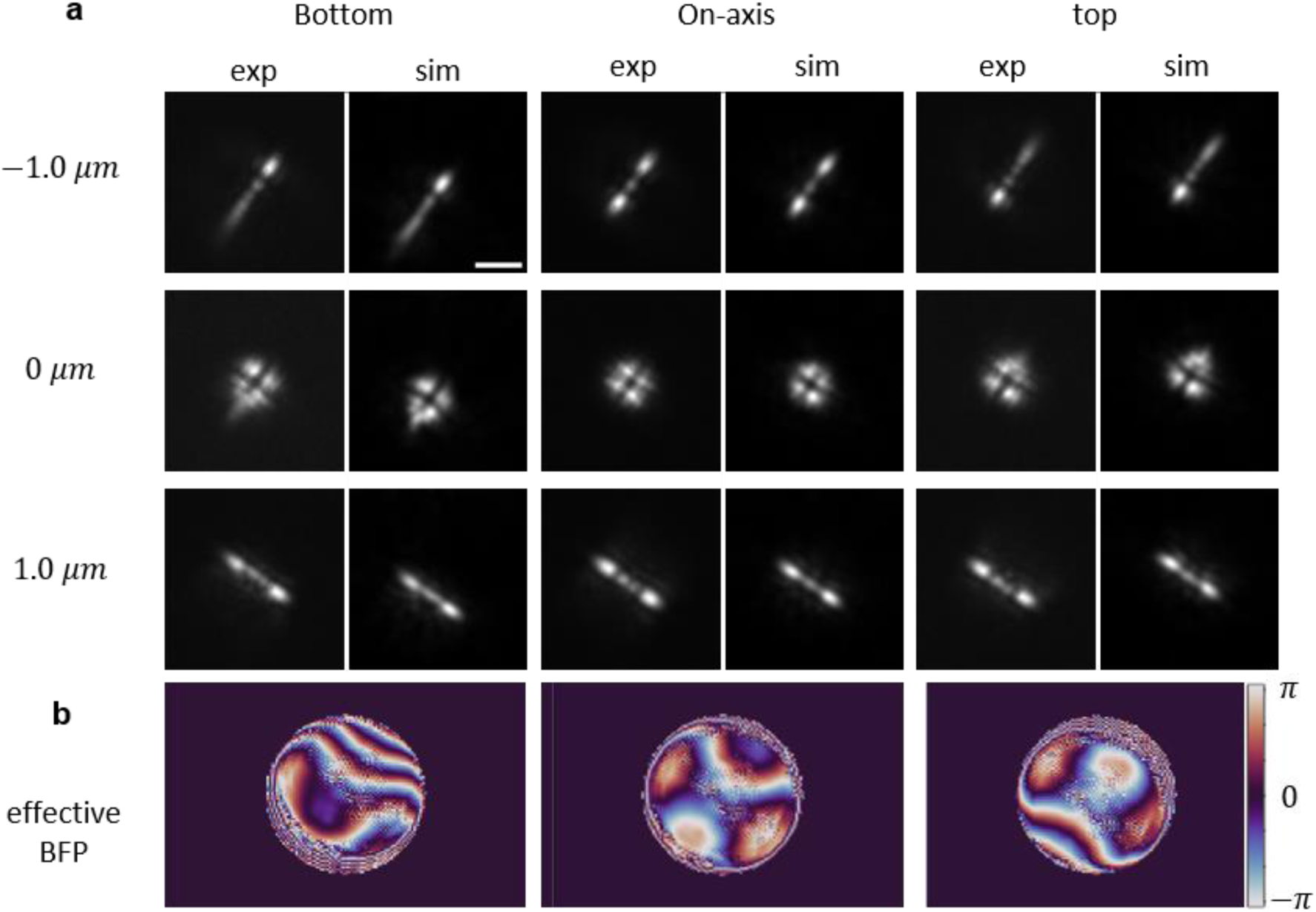
Phase retrieval results for displaced DOE. **a**, Comparison of experimental PSFs (“exp”) and forward-model simulations (“sim”) at three defocus positions (−1, 0 and +1 μm). Simulations were generated using the phase mask and displacement parameter recovered by displaced-mask phase retrieval. **b**, Effective pupil-plane phase modulation corresponding to the recovered displaced-mask model for the three emitter field positions. Because the beam shifts at the displaced mask plane according to emitter’s lateral displacement, emitters at different field positions experience different effective phase modulations, resulting in field-dependent PSFs. Left column: emitter in the bottom region of the field of view, pixel coordinates (368,826). Middle column: on-axis emitter, pixel coordinates (477,573). Right column: emitter in the top region of the field of view, pixel coordinates (582,358). The optimized mask displacement was 27 mm. Scale bar: 2 µm.

For numerical stability and physical plausibility, mild regularization or smoothing could be applied to the phase mask during optimization in future work. This will suppress unrealistically high-spatial-frequency structure in the retrieved mask while preserving the dominant modulation required to reproduce the measured PSFs. The final recovered mask was then used throughout the remainder of the pipeline as the optical prior of the system.

### Supplementary Note 8 – Detection-threshold sensitivity and axial-profile analysis

The reconstruction, visualization and FRC procedures are described in the Online Methods. Here, we further examine how the network-output threshold affects localization density and apparent axial profile width in the field-dependent reconstruction.

The reconstructions presented in the main text were generated using the same network-output intensity threshold of 30, for both the standard AutoDS3D and field-dependent analyses. This common threshold provides a direct comparison between the two workflows without manually tuning the detection threshold for each method. Under this condition, the field-dependent reconstruction produced more localizations over the full field of view than the standard AutoDS3D reconstruction, while the number of detected localizations near the optical axis remained similar. This is consistent with the expected role of the field-dependent model, which improves recovery of emitters away from the optical axis, where shift-invariant calibration is less accurate.

We further examined the effect of increasing the field-dependent reconstruction threshold from 30 to 50, on the reconstruction near the optical-axis. This higher threshold reduced the number of accepted localizations but narrowed the axial profiles extracted from the representative ROIs shown in Supplementary **Fig. S8**. In the two ROIs shown (similar to the ROIs in Fig 2f and h in the main text), the standard AutoDS3D reconstruction at threshold 30 yielded axial FWHM values of approximately 92 nm and 93 nm. The field-dependent reconstruction at intensity threshold 30 yielded broader axial profiles of approximately 136 nm and 115 nm, whereas increasing the field-dependent threshold to 50 reduced these values to approximately 92 nm and 88 nm, respectively.

**Fig. S8:**
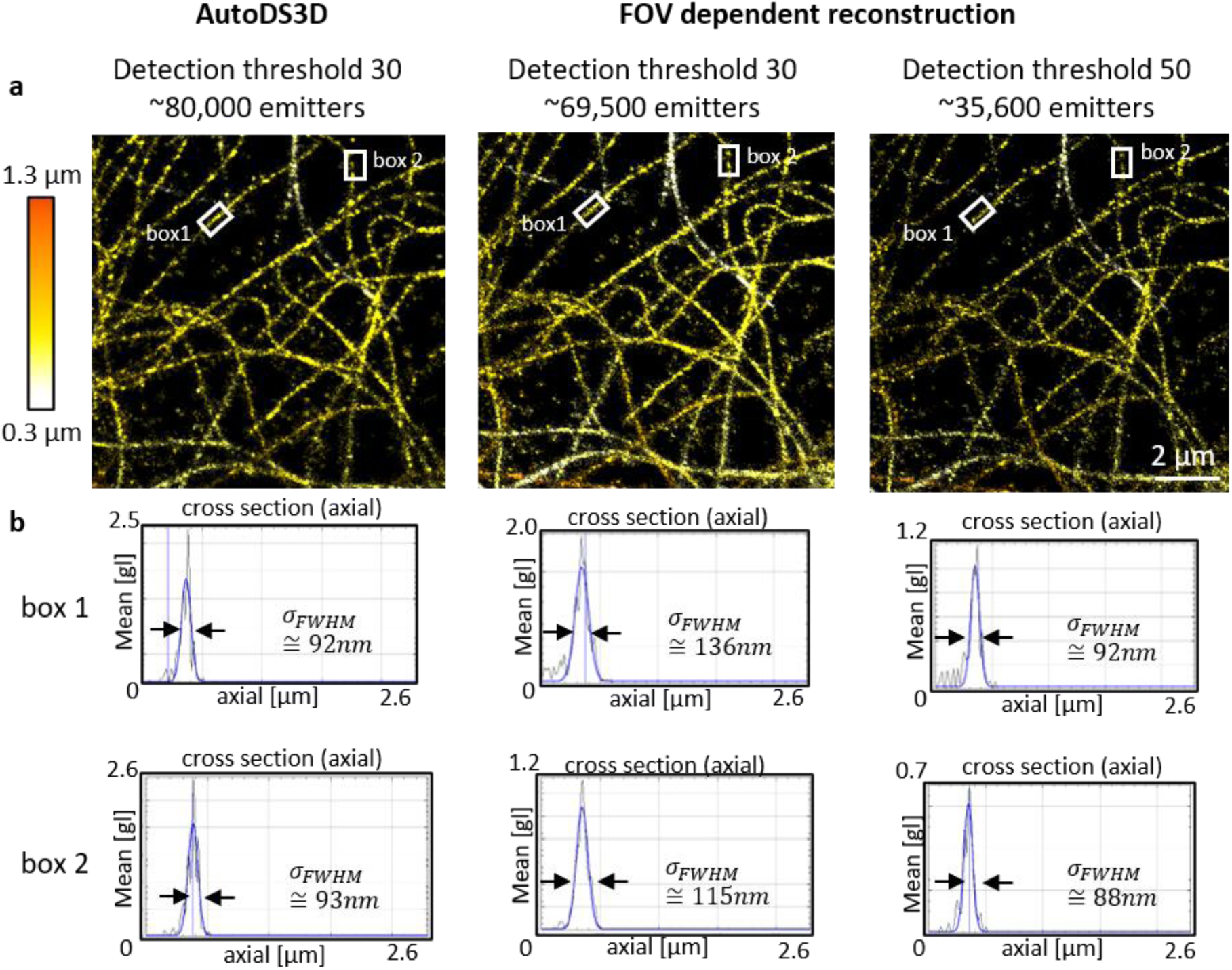
Effect of detection threshold on axial profile width. **a,** 3D STORM reconstructions of microtubules comparing standard AutoDS3D at threshold 30, FOV-dependent reconstruction at threshold 30 and FOV-dependent reconstruction at threshold 50. The same field of view is shown in all panels near the optical axis, with two representative regions of interest marked by white boxes. Color indicates axial position. **b,** Axial cross-sectional profiles measured from the marked ROIs. At the common threshold of 30, the FOV-dependent reconstruction produces broader axial profiles than AutoDS3D in these near-axis regions. Increasing the FOV-dependent threshold to 50 narrows the axial profiles to values comparable to AutoDS3D, but reduces the number of accepted localizations. These results illustrate the trade-off between localization density and apparent axial width in the current FOV-dependent reconstruction.

These results highlight a trade-off of the current PnP reconstruction workflow. Direct mask insertion simplifies the optical implementation but produces field-dependent PSFs, shifting part of the burden to computation. At lower threshold, the field-dependent reconstruction recovers more emitters across the full field of view, including regions where a shift-invariant model fails, but lower-confidence detections can broaden local axial profiles, resulting in an advantage for the standard AutoDS3D near the optical-axis. At higher threshold, the reconstruction becomes more selective and yields sharper axial profiles, but discards a substantial fraction of localizations, including near the optical axis. Future implementations may benefit from improved confidence calibration, spatially adaptive thresholding and reconstruction algorithms tailored to displaced- mask PSFs.

### Supplementary Note 9 – Microscope adapters and alternative mounting geometries

The PnP component demonstrated in this work was implemented on a Nikon Ti2 microscope using an adapter designed to fit the existing DIC slot (**Fig. S9**a). This slot-based design allows the phase mask to be inserted into the microscope infinity space without modifying the optical path or moving the camera.

However, we also designed an Olympus-compatible implementation. In this design, a circular 3D- printed phase-mask holder is attached magnetically to an existing Olympus DIC-slot adapter (**Fig. S9**b). This modular geometry allows a single microscope adapter to be reused with different phase-mask holders, while the magnets provide repeatable mechanical positioning and simplify mask exchange.

We also considered a more general objective-mounted configuration, in which the phase mask is held in a threaded adapter attached directly below the objective lens (**Fig. S9**c). A similar mounting concept was previously used for an extended-depth-of-field phase mask by Opatovski et al.^5^. Although this configuration is not strictly PnP in the slot-based sense, because the mask is mounted at the objective rather than inserted into an existing microscope slot, it provides a simple alternative for microscopes without accessible DIC or filter-slot positions. Together, these examples illustrate that the PnP-PSFE concept can be implemented using different mechanical interfaces, depending on the available access points of the microscope.

**Fig. S9.**
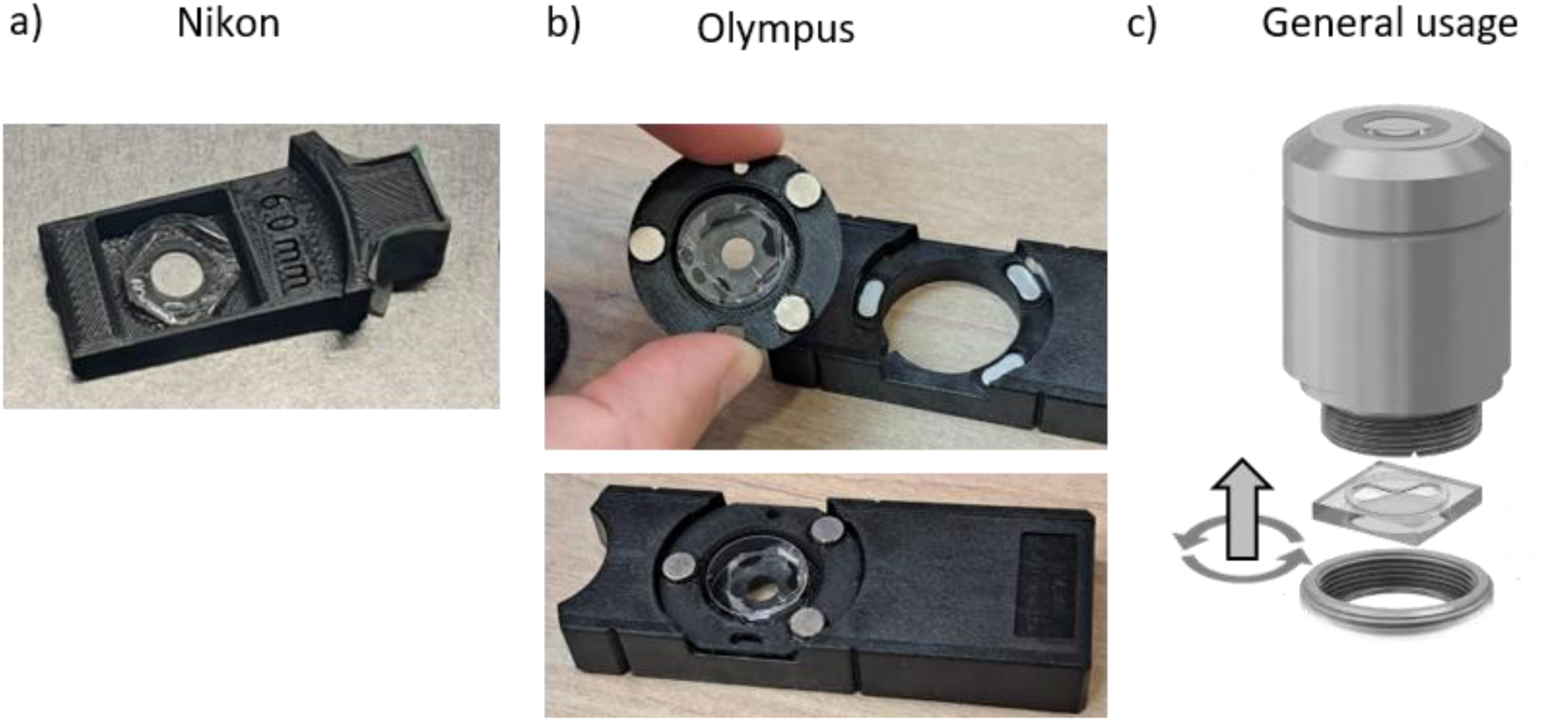
Microscope adapters for PnP phase-mask integration. **a,** Nikon adapter designed according to the geometry of Nikon DIC components. The phase mask is mounted in the adapter and inserted into the existing microscope slot. **b,** Olympus-compatible modular adapter. The circular 3D-printed phase-mask holder contains the mounted phase mask and attaches to an Olympus DIC-slot adapter using magnets, allowing different phase masks to be exchanged while using the same microscope adapter. **c,** Objective-mounted configuration, in which the phase mask is held in a threaded adapter attached directly below the objective lens. This geometry does not preserve the same slot-based plug-and-play operation but provides a simple alternative for microscopes without a suitable insertion slot.

## Notes

https://doi.org/10.5281/zenodo.21648990

https://doi.org/10.5281/zenodo.21648914

https://doi.org/10.5281/zenodo.21648812

### Competing Interest Statement

The authors have declared no competing interest.

### Summary of Updates

This version of the manuscript has been revised to update the Author list and Acknowledgments section

## References

1. Lelek, M. et al. Single-molecule localization microscopy. Nat Rev Methods Primers 1, 39 (2021).

2. Rust, M. J., Bates, M. & Zhuang, X. Sub-diffraction-limit imaging by stochastic optical reconstruction microscopy (STORM). Nat Methods 3, 793–796 (2006).

3. Betzig, E. et al. Imaging Intracellular Fluorescent Proteins at Nanometer Resolution. Science 313, 1642– 1645 (2006).

4. Hess, S. T., Girirajan, T. P. K. & Mason, M. D. Ultra-high resolution imaging by fluorescence photoactivation localization microscopy. Biophys J 91, 4258–4272 (2006).

5. Huang, B., Wang, W., Bates, M. & Zhuang, X. Three-dimensional super-resolution imaging by stochastic optical reconstruction microscopy. Science 319, 810–813 (2008).

6. Pavani, S. R. P. et al. Three-dimensional, single-molecule fluorescence imaging beyond the diffraction limit by using a double-helix point spread function. Proc Natl Acad Sci U S A 106, 2995–2999 (2009).

7. Nehme, E. et al. DeepSTORM3D: dense 3D localization microscopy and PSF design by deep learning. Nat Methods 17, 734–740 (2020).

8. Gahlmann, A. et al. Quantitative Multicolor Subdiffraction Imaging of Bacterial Protein Ultrastructures in Three Dimensions. Nano Lett. 13, 987–993 (2013).

9. Shechtman, Y., Weiss, L. E., Backer, A. S., Lee, M. Y. & Moerner, W. E. Multicolour localization microscopy by point-spread-function engineering. Nature Photon 10, 590–594 (2016).

10. Shechtman, Y., Sahl, S. J., Backer, A. S. & Moerner, W. E. Optimal Point Spread Function Design for 3D Imaging. Phys. Rev. Lett. 113, 133902 (2014).

11. Orange kedem, R. et al. Near index matching enables solid diffractive optical element fabrication via additive manufacturing. Light Sci Appl 12, 222 (2023).

12. Saguy, A. et al. One-click reconstruction in single-molecule localization microscopy via experimental parameter-aware deep learning. npj Imaging 3, 61 (2025).

13. Ferdman, B., Saguy, A., Xiao, D. & Shechtman, Y. Diffractive optical system design by cascaded propagation. *Opt. Express*, OE 30, 27509–27530 (2022).

14. Xiao, D. et al. Large-FOV 3D localization microscopy by spatially variant point spread function generation. Science Advances 10, eadj3656 (2024).

15. Cohen, O. R. et al. Compact Spectral Encoding Microscopy by Terrace Grating Optics. ACS Photonics 13, 1407–1416 (2026).

16. Joshi, P. et al. High-speed volumetric single-molecule imaging using dual-wavelength light sheets and PSF-engineered enhanced biplane detection. 2026.06.20.733419 Preprint at 10.64898/2026.06.20.733419 (2026).

17. Saliba, N., Cheng, S., Joshi, P. & Gustavsson, A.-K. Versatile and Scalable Reflective Micromirrors for Single-Objective Light Sheet Microscopy. Nano Lett. 10.1021/acs.nanolett.6c01709 (2026)

## Supplementary References

1. Orange Kedem, R. et al. Near index matching enables solid diffractive optical element fabrication via additive manufacturing. Light Sci Appl 12, 222 (2023).

2. Cohen, O. R. et al. Compact Spectral Encoding Microscopy by Terrace Grating Optics. ACS Photonics 13, 1407–1416 (2026).

3. Saleh, B. E. A. Fundamentals of Photonics I: Optics/ Bahaa E.A. Saleh, Malvin Carl Teich. (Wiley, Hoboken, NJ, 2019).

4. Petrov, P. N., Shechtman, Y. & Moerner, W. E. Measurement-based estimation of global pupil functions in 3D localization microscopy. Opt. Express, OE 25, 7945–7959 (2017).

5. Opatovski, N. et al. Depth-enhanced high-throughput microscopy by compact PSF engineering. Nat Commun 15, 4861 (2024).

